# Why multiple infections favour virulent parasites

**DOI:** 10.1101/258004

**Authors:** Mircea T. Sofonea, Samuel Alizon, Yannis Michalakis

## Abstract

It is now a fact that several strains/species (hereafter types) of parasites circulate in natural host populations. Parasite polymorphism can even occur within the same host, where distinct parasite types can interact in various ways. This can affect their transmission and, therefore, their evolution. We still lack general predictions regarding the evolution, in such multiple infection contexts, of virulence – the infection-induced host mortality, essentially because its emanation from within-host growth was often ignored so far. Here, we explicitly investigate within-host interactions, within-host competition outcomes, epidemiological dynamics and evolutionary invasibility using a formalism as general as possible. Focusing on chronic dimorphic infections caused by horizontally-transmitted microparasites, we apply both dynamical systems and probabilistic approaches to this bottom-up sequence of dynamics to explore the evolutionary outcomes. We show that within-host growth traits are under strong selective pressure and when small mutations affect them, most of the surviving mutants are more virulent than their resident. We thus identify a robust and unavoidable selection bias towards higher virulence.

## 1 Introduction

Virulence has been an intriguing paradox since parasites that reduce the lifetime of their host concomitantly reduce their opportunities to be transmitted and thus one would expect them to be counter-selected (Smith, 1887; Méthot, 2012). Adaptive evolution of virulence has been reviewed several times (Dieckmann, 2002; Alizon and Michalakis, 2015; Cressler et al., 2015) and one can distinguish three main reasons to why parasites may harm their hosts: 1) virulence trades off with other features of the parasite‘s life cycle (transmission, recovery, free stage persistence) (Anderson and May, 1982; Gandon, 1998; Alizon, 2008; Alizon et al., 2009; Alizon and Michalakis, 2011), 2) virulence is associated with within-host competitiveness (Alizon et al., 2013) and 3) virulence is the by-product of ‘short-sighted’ within-host evolution (Levin and Bull, 1994). Notice that each of these explanations is self-contained but they are not mutually exclusive.

The fact that virulence may be driven by the competition between parasites seems relevant as it is now well established that a diversity of parasite types (species, strains, genotypes) share the same host population (Petney and Andrews, 1998; Lord et al., 1999; Juliano et al., 2010; Rigaud et al., 2010; Balmer and Tanner, 2011). When several types of parasites circulate in the same host population, they not only compete for susceptible hosts to infect but they also compete within the host for limiting resources (Mideo, 2009). When simultaneously inside the same host, parasite interactions, may they be direct or indirect, positive or negative, within the same type or between types (Bashey, 2015), can make coexistence impossible, as in any ecological community (Barabás et al., 2016). This subsequent diversity of within-host outcomes (Sofonea et al., 2017) amplifies the richness of the epidemiological dynamics governed by transmission, recovery and death of the thus numerous classes of hosts (Sofonea et al., 2015; Kucharski et al., 2016). Besides, within-host coexistence of several parasite types can modulate the rate at which a host dies from the multiple infection with respect to single infections (Griffiths et al., 2011). It follows that the virulence undergone by a multiply infected host has to be distinguished from the virulence of a parasite, usually revealed in single infections (de Roode et al., 2005).

The first evolutionary epidemiological models accounting for parasite polymorphism exposed an evolutionary increase of virulence comparable to the selfish tendency to over-exploit a common good, as in any tragedy of the commons (Levin and Pimentel, 1981; Bremermann and Pickering, 1983; Nowak and May, 1994; van Baalen and Sabelis, 1995; May and Nowak, 1995; Frank, 1996). The generality of this conclusion has however been challenged by later models that assumed host exploitation to rely on a collective action (such as public good production or parasite load limitation) which tends to decrease parasite virulence (Brown, 1999; Brown et al., 2002; West and Buckling, 2003). These results were in turn contradicted when accounting for trade-offs and epidemiological feedbacks (Alizon and Lion, 2011). This in the end reveals that, up to now, theoretical research on the consequences of multiple infections has failed to provide a unified answer to the selective pressure parasite virulence undergoes. This is at least partly explained by the tendency to increase the complexity of the model from the top (Alizon, 2013a) while keeping within-host dynamics, and therefore how virulence arises, unaddressed (but see Alizon and van Baalen (2005)).

Because of their challenging analysis, the within-host dynamics have been ignored or oversimplified by an impressive majority of evolutionary epidemiological models involving at least two parasite types (Alizon, 2013a; Cressler et al., 2015). More precisely, most of these models arbitrarily forced all growth outcomes inside multiply inoculated hosts to result either to the dominance of a unique type (Levin and Pimentel, 1981) or to the coexistence of all inoculated types (Bremermann and Pickering, 1983). The first case is known as superinfection and corresponds to the fast competitive exclusion of all parasite types that are less competitive than the surviving parasite type. Competitiveness is almost always assumed to be positively correlated with virulence, which is both limiting and arguable (Sofonea et al., 2017). The second case is known as coinfection and corresponds to stable (or meta-stable) coexistence between all inoculated parasite types the transmission rates of which are often assumed to be identical to that of single infections, which is also limiting and arguable (Sofonea et al., 2017). Although Nowak and May acknowledged these two cases might be biologically extreme, they claimed “to bracket the reality of polymorphic parasites” (May and Nowak, 1995). It eventually became common in the literature that the epidemiology of any biological system with multiple infections lies somewhere in a continuum between super and coinfection e.g. (Mosquera and Adler, 1998; Boldin and Diekmann, 2008; Alizon, 2013a). Our previous work has shown, however, that explicitly considering within-host dynamics reveals a diversity of within-host outcomes (Sofonea et al., 2017), questioning the relevance of imposing within-host outcomes when investigating the evolution of parasite traits.

Finding a general trend in virulence evolution with multiple infections thus requires constructing a new framework based on the parasite growth traits (Cressler et al., 2015). By means of bottom-up emergence and dynamical nesting, this approach should be able to capture, and weight, a diversity of multiple infection scenarios so far separated or ignored. In a previous work (Sofonea et al., 2017), we introduced and detailed, both formally and biologically, the notion of infection pattern which captures the within-host outcomes of all sets of parasite types. Intuitively, an infection pattern is the way each parasite type behaves – *i.e.* either survives or vanishes – in presence of any combination of other parasite types (including when the focal type is alone).

First, this notion generalizes the super/coinfection dichotomy. We indeed showed that even with only two parasites, there can be more than a hundred of these patterns. Interestingly, several of the patterns we uncovered theoretically have been known in the empirical literature for decades and cannot be modelled as any mixture of super and coinfection. This is for example the case when the first inoculated type always outcompetes the second one, as for the mutual exclusion of phages in *E. coli* (Delbrück, 1945), a pattern we referred to as priorinfection (Sofonea et al., 2017). Second, infection patterns allows us to root the epidemiological network and transmission routes in the within-host dynamics while relaxing the assumption that links competitiveness to virulence.

In the present article we take advantage of the previously introduced infection pattern typology for dimorphic infections (Sofonea et al., 2017) to investigate how mutations on parasite growth traits affect the average virulence of the parasite meta-population (*i.e.* at the between-host level) through nesting a classical two-species competitive Lotka-Volterra model for within-host dynamics into into an extended version of the Susceptible-Infected-Susceptible (*SIS*) model developed in (Sofonea et al., 2015). The integrated principle of this bottom-up approach is exposed in Figure 1. We finally interpret the results as an evolutionary force that acts on parasite virulence, examining the effects of both small and large mutations.

**Figure 1:**
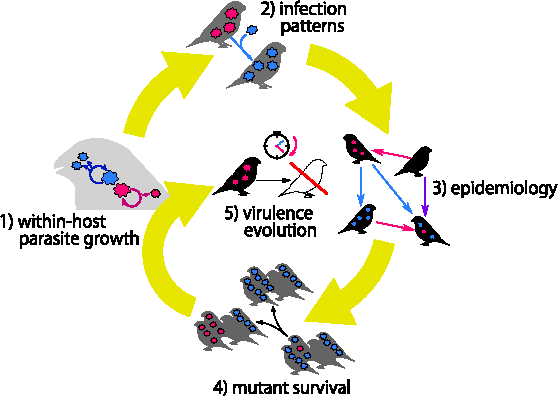
Overview of the bottom-up framework used to address virulence evolution. This figure illustrates the general approach used in this work that involves three dynamical levels: the within-host dynamics (which also includes secreted compounds kinetics) (step 1), the between-host dynamics (step 3) and the evolutionary dynamics (step 5). Each of these three levels is characterized by a proper model respectively inspired from the competitive Lotka-Volterra equations, the *SIS* model and general probability theory. They are nested one into the other by two key theoretical linkages: the infection pattern typology (step 2) projects the within-host growth outcomes into epidemiology while mutant survival (step 4) deducts from the epidemiology the result on virulence evolution, the goal of our study viewed through the prism of parasite growth traits (hence the last arrow‘s detour). See references in main text.

We provide in the appendix the full description and analysis of the models (Supplementary Methods part) and some more technical results (Supplementary Results part).

## 2 The model

### 2.1 Within-host dynamics

The parasite population dynamics that take place within a host are assumed to be governed by the interactions between parasites individuals. After inoculation, the growth of each parasite type population depends on the parasite load of both types but also on the concentrations of secreted compounds in the medium that can originate from the parasites or the hosts. We refer to compounds that positively affect parasite growth as public goods, *e.g.* siderophores for bacteria (West and Buckling, 2003) or replication enzymes for viruses (Huang and Baltimore, 1970). Those with a negative effect are either parasite spite such as bacteriocins (Gardner et al., 2004) in bacteria and entry receptor down regulators (Nethe et al., 2005) in viruses, or host defense molecules induced by the infection, *e.g.* lactoferrin and siderocalin (Skaar, 2010). We thus do not regard within-host parasite growth as a simple exploitative competition but we allow for interference competition, apparent competition and cooperation as well (Read and Taylor, 2001; Cressler et al., 2015).

These diffusible compounds are assumed to be renewable, which is the case of any enzyme and known for siderophores (Neilands, 1995), lactoferrin (Farnaud and Evans, 2003), some bacteriocins (Riley and Chavan, 2007). They are also assumed to be mostly removed from the medium through parasite unrelated processes such as self-denaturation, dilution or host immunity. If one of these two conditions is met, a time scale separation between compound kinetics and parasite replication holds and the parasite within-host dynamics can be reduced to a classical competitive Lotka-Volterra model (Lotka, 1925; Volterra, 1928) illustrated in Figure 2 for the dimorphic case to which we restrict our approach (the full model and the proof of the reduction are provided appendix B.1). Labelling the parasite types 1 and 2, the within-host dynamics only depends on six parameters: the intrinsic growth rates, *ϱ*_1_, *ϱ*_2_, that quantify the speed at which each parasite individual would reproduce if completely isolated, the self interaction traits, *m*_1,1_, *m*_2,2_, that quantify the strength of intra-type competition, and the cross interaction traits, *m*_1,2_, *m*_2,1_, that quantify the strength of inter-type competition. Note that from the point of view of a given type, let us say 1, both *m*_1,1_ and *m*_1,2_ capture the competitive pressure experienced by 1 while *m*_2,1_ captures its competitiveness. These interpretations can likewise be given in cooperation terms.

**Figure 2:**
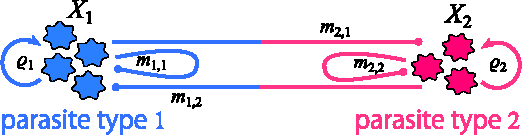
Within-host model. The load of parasite type 1 (*X*_1_) varies according to its instantaneous growth rate, which is the sum of its intrinsic growth rate *ϱ*_1_, intra-type interaction effect *m*_1,1_ *X*_1_ and inter-type interaction effect *m*_2,1_ *X*_2_, where *X*_2_ denotes the load of parasite type 2. The same holds symmetrically for parasite type 2, but notice that the cross interaction traits are not necessarily symmetric (*m*_1,2_ *[negationslash]*= *m*_2,1_), even though they are shared traits (as signified by their colors). More generally, the four interaction traits *m_i,j_* capture density-dependent, beneficial and detrimental compounds and host environment mediation effects.

The within-host system possesses four equilibria (appendix B.2) – one uninfected, two singly infected and one doubly infected. Each of them corresponds to a host class, which we define as the set of parasite types that chronically infect a host. Besides recovery addressed below, hosts may change class owing to inoculation challenges.

In order to assess the consequence of an inoculation, we investigate the feasibility and the stability of each fixed point given the inoculated parasite types (see appendix B.3 for details). Figure 3 shows the within-host growth of the two parasite types when sequentially inoculated in the same host, according to four different parameter sets. As already pointed in Sofonea et al. (2017), these results illustrate that our simple within-host model can generate a diversity of within-host outcomes such as type replacement (**A**), stationary coexistence (**B**), infection burst (**C**) and mutual exclusion (**D**). Notice that only the first two cases are addressed by the super/co-infection dichotomy (Nowak and Sigmund, 2002).

**Figure 3:**
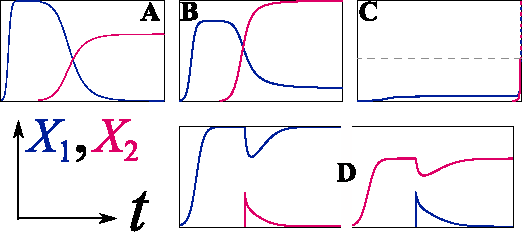
Examples of inoculation outcomes. Four representative outcomes generated by the within host model are shown in terms of parasite loads through time, (*X*_1_, *X*_2_) (*t*). In all panels, parasite type 1 is first inoculated alone in the host and its load reaches a plateau, the value of which is equal in the four parameter sets for the sake of comparison). Few time units later, the second parasite type is inoculated. **A**) Type replacement: parasite type 2 grows and outcompetes parasite type 1 (notice that it reaches a lower plateau but this is not necessary). **B**) Stationary coexistence: both parasite types reach stationarity (notice that parasite type 1 then reaches another plateau). **C**) Infection burst: once together, the two parasite types’ growth explode and therefore exceeds the load threshold above which an acute host immune response is triggered (here arbitrarily set to ten times the stationary parasite load of parasite type 1 and denoted by the dashed line). **D**) Mutual exclusion, even inoculated at high dose, parasite type 2 fails to grow and rapidly vanishes while parasite type 1 recovers its plateau (left sub-panel). Inverting the inoculation order shows the opposite outcome (right sub-panel). The parameter sets used for numerical integration are given in appendix B.1.

### 2.2 Infection patterns

The space of within-host traits can be split into regions that share the same within-host outcomes for each inoculation challenge. These regions are called infection patterns and have been thoroughly discussed in Sofonea et al. (2017), both abstractly (see the mapping formalism in the appendix therein) and biologically (hence the associated terminology and biological examples). The present model generates seven non trivial infection patterns illustrated in Figure 4 (see proof in appendix C). These patterns have already been exposed in Sofonea et al. (2017), where biological support and references are provided. Besides the aforementioned superinfection and coinfection patterns, we emphasize on priorinfection, where the two kinds of single infections are possible but none of the parasite types can take over a host already infected by the other, and latinfection, where one parasite type is absent from all within-host outcomes (although it can be transiently observed).

**Figure 4:**
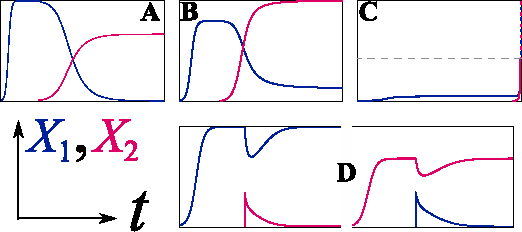
Infection patterns. Graphical representation and names of the seven non trivial infection patterns generated by the within-host model. We only provide one example of asymmetric patterns for which a distinct twin can be obtained by swapping the type labels and their name is followed by 1. Circled labels represent host classes and arrows transitions between host classes through a specified inoculation. Full blue arrows represent inocula of type 1 alone, full red arrows represent inocula of type 2 alone, full purple arrows represent inocula of both types, blue and purple dashed arrows represent inocula of type 1 and possibly type 2, and red and purple dashed arrows represent inocula of type 2 and possibly type 1. For the sake of clarity, inoculation challenges that do not produce any host class transition are not shown. (More details on graphical conventions and naming of infection patterns are available in Sofonea et al. (2017).)

It is worth noticing moreover that, unlike previous models that assume that the most virulent type can take over any infected host (Levin and Pimentel, 1981; Nowak and May, 1994), our model is more general and allows competitive exclusion by the less virulent type, provided that virulence is an increasing function of parasite load (see appendix D.3), as empirical studies on various host-parasite systems suggest (Ebert and Mangin, 1997; Kover and Schaal, 2002; de Roode et al., 2008; Fraser et al., 2014; Sy et al., 2014). More generally, this reminds that assuming a unique infection pattern while investigating the space of within-host growth traits can lead to erroneous interpretations.

### 2.3 Between-host dynamics

Accounting for parasite polymorphism in epidemiology is a challenge both in terms of modeling and biological interpretation (Metcalf et al., 2015). Indeed, the number of host compartments increases exponentially with the number of parasite types if all type combinations are allowed to coinfect (Sofonea et al., 2015; Kucharski et al., 2016). For the sake of both biological intelligibility and mathematical tractability, we adapt the classical Susceptible-Infected-Susceptible (*SIS*) model (Keeling and Rohani, 2008) to parasite dimorphism. We however endow this model with very general features.

Here, the functions that govern host demography, with respect to host densities, and epidemiological rates (transmission, virulence and recovery), with respect to parasite load, only have to satisfy minor and common assumptions (*e.g.* transmission does not decrease with parasite load). Importantly, these functions are otherwise left unspecified, that is we do not assign them any analytical expression. We emphasize on the fact that this guarantees a great generality of the subsequent results which will therefore be valid for a diversity of demographic dynamics and physiological responses hence also preventing them from structural sensitivity (Wood and Thomas, 1999; Fussmann and Blasius, 2005).

Besides, we allow the doubly infected hosts to transmit the three possible kinds of inocula (1 alone, 2 alone and 1 & 2) and, in addition to recovery, all compartment transitions are possible so that the model captures all infection patterns previously introduced. The flow diagram and more information about the model are given in Figure 5. The associated equations and the analysis of the model are provided in appendix D.

**Figure 5:**
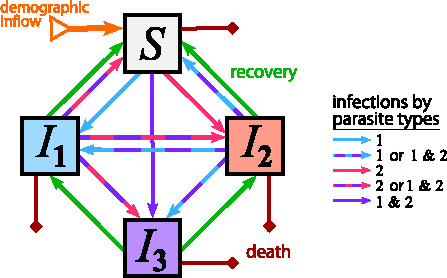
Flow diagram of the generic *SIS* model with parasite dimorphism. The between-host dynamics characterize the density of four host compartments (boxes) through time, namely the susceptible hosts *S*, the hosts singly infected by parasite type 1 *I***_1_**, the hosts singly infected by parasite type 2 *I***_2_** and the doubly infected hosts. Parasite transmission is assumed to be purely horizontal so all new hosts (offspring or immigrants) are susceptible (orange arrow). Hosts change compartment either through infection (blue, pink, purple and mixed arrows) or through recovery (green arrows). Infections result from transmission of one or both parasite types from both singly or doubly infected hosts. Recovery is possible one parasite at a time. Finally, mortality (deep red arrows) affects all host compartments but at different rates: natural mortality is shared but virulence, that is additional mortality due to infection, can differ among infected host compartments.

### 2.4 Mutational setting

Virulence originates from the parasite ability to grow within a host. It can therefore be seen through the prism of stationary parasite load (as, for instance, in Fraser et al. (2014)), which is the combination of two within-host growth traits: intrinsic growth rate and self interaction.

Let us consider that a monomorphic parasite is already circulating in a host population (not necessarily at epidemiological equilibrium) and, among its offspring, one mutant individual possesses traits that differ from those of its resident ancestor. Regarding the ancestor and its mutant as respectively parasite types 1 and 2, we write *ϱ*_2_ = *ϱ*_1_ + *R* and *m_i,j_* = *m*_1,1_ + *M_i,j_* where *R* and *M_i,j_* are random variables. The greatest standard deviation of these random variables hereafter serves as a proxy for the overall mutational effect.

In the following, we focus on mutants that differ only slightly from their ancestor as it allows us to fully characterize its survival in spite of the model‘s generality, while being a usual assumption in theoretical evolutionary biology (Taylor, 1989; Geritz et al., 1998). To do so, we examine a diversity of mutational settings we distinguish according to two complementary properties: the mutational distribution – that governs the spreading shape of the probabilities with which the mutant traits take their values (*i.e.* the laws of *R* and *M_i,j_*) – and the mutational scenario – that governs the dominance and correlations between mutant traits. We hereafter verbally expose a simplified view of these settings but see appendix E for their mathematical formulation.

In order to capture a large range of mutational processes, we consider three different mutational distributions, a) the fixed step distribution, *i.e.* mutations that, in first approximation, add or subtract with equal probabilities the same quantity to the ancestor trait value, b) the Gaussian distribution, where the mutant trait values are normally distributed around those of the ancestor (Wolf et al., 2007), and c) the generic distribution, that is any probability distribution symmetrically distributed around the ancestor trait values. The fixed step and generic mutational distributions are treated analytically while the Gaussian mutational distribution is used for numerical confirmations (and larger mutation effects, see below).

As for the mutational scenarios, we consider the cases where #1 - the mutation results in a change in intrinsic growth rate, #2 - the mutation mainly lies in the interaction traits, #3 - the mutation independently affects all four traits simultaneously in comparable absolute value (we call it the unconstrained, or free, scenario), #4 - the traits are strongly traded-off, e.g. an increase in intrinsic growth rates necessarily increases the competitive pressure undergone by the mutant and lowers its competitiveness.

We can now implement these mutational settings into the previously developed bottom-up ap-13 proach, from within-host interactions to epidemiological dynamics and investigate the proportion of mutants that survive with respect to their virulence, which is the support of virulence evolution. The endpoint results of this analysis are the proportions of surviving mutants that are more virulent than their ancestor obtained under all combinations of mutational distributions and mutational scenarios (see subsection 3.1 below).

For the sake of completeness, we also extend our investigations to large mutation effects. Because they raise analytical difficulties, we limit ourselves to numerical simulations of only one, but representative, case: the Gaussian mutational distribution under the free mutation scenario (#3) (see subsection 3.2 below).

## 3 Results

### 3.1 Small mutations

If mutations have a small phenotypic effect, it is easy to show that under any of the mutation settings described above, exactly one half of the generated mutants are more virulent than their ancestor. Besides, the only infection patterns that emerge are superinfections, coinfection and priorinfection. Following the next-generation method (van den Driessche and Watmough, 2002; Hurford et al., 2010), we derive the invasion fitness of the mutant the sign of which is, at least partially, governed by the infection pattern. Hence, we show that a mutant can survive, in spite of its ancestor‘s anteriority in the host population, only when, within one doubly inoculated host, it always excludes its ancestor (superinfection 2), coexists with it (coinfection) or when the the mutant and its ancestor exclude each other according to the anteriority inside the host. More precisely, all mutants that establish a super-infection 2 or a coinfection survive whatever their virulence. In contrast, the survival of the mutants involved in a priorinfection depends on virulence, as the sign of the invasion fitness changes according to the shape of the epidemiological trade-offs that govern their capacity to spread among hosts, usually quantified by the basic reproduction number *R*_0_(Diekmann et al., 1990). Consequently, under prior-infection, the mutants that survive are either those that are more virulent than their ancestor; if *R*_0_ increases with virulence, or, those less virulent if *R*_0_ decreases with virulence. Investigating whether R0 increases or decreases with virulence is exactly equivalent to investigating whether the resident‘s virulence is located on an ascending or a descending slope of the unidimensional *R*_0_ landscape with respect to virulence. Since we do not investigate any particular mutant nor any particular *R*_0_ landscape we make the neutral assumption that the direction of the slope on which the resident‘s virulence is located has as equal chances to be ascending or descending. Hence, because half of the mutations increase virulence, it results that half of the mutants involved in priorinfection survive in the end. If a mutant establishes any other pattern (essentially superinfection 1, but this holds for latinfection 2 and ultrainfection as well), it always goes extinct. The results about how infection patterns affect the mutant‘s probability of survival are summarized in Table 1.

**Table 1:** Mutant survival according to infection pattern.

| infection pattern | superinfection 2 | coinfection | priorinfection | other patterns |
| --- | --- | --- | --- | --- |
| probability that the mutant survives given the infection pattern | 1 | 1 | $1/2$ | 0 |

Intuitively, the mutants that possess both properties of higher virulence and survival are a minority. However, when we focus on the proportion of more virulent mutants among the surviving mutants, by combining results from Table 1, the frequency of each infection pattern and their statistical association with virulence (see appendix F), we find that more virulent mutants always constitute the majority of surviving mutants. This observed selection bias towards higher virulence is supported by all combinations of investigated mutational distributions and mutational scenarios, as shown in Figure 6, where all analytical values or numerical estimates of the proportion of more virulent mutants among surviving mutants lie in the right half side of the gradient.

**Figure 6:**
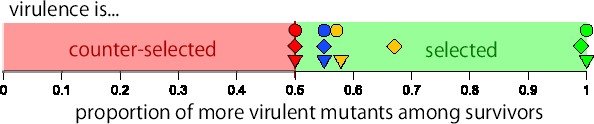
Virulence selection bias in several mutational settings. The horizontal position of the triangles, circles and rhombi gives the proportion of surviving mutants that are more virulent than their ancestor when the mutational distribution is a fixed step, generic and Gaussian respectively. The results are analytical for the first two distributions and numerical for the third (10^5^ simulations for each mutational setting). Colors denote the mutational scenario: fully constrained in red, competition-driven in blue, unconstrained in orange and growth-driven in green Since the solution in the generic distribution of unconstrained mutation is not unique, only a lower boundary is shown. The effect of the epidemiological trade-off is neutralized by averaging over a distribution of resident virulence assumed symmetric with respect to the epidemiologically optimal virulence.

Although the precise results rely on tedious calculations (exposed in appendix F), it is possible to qualitatively approach this apparently robust virulence selection bias by means of a marginal simplification of the mutational setting and elementary geometry, as provided in Figure 7. A simplified interpretation of the results would state that, on one hand, a higher virulence is achieved through faster growth and greater self cooperation compared to the respective ancestor‘s traits. On the other hand, a mutant survives through faster growth and competition escape. Most mutants that grow faster are also more virulent, while those who escape their ancestor‘s competition can nonetheless present a greater self cooperation which makes some of them more virulent as well. In the end, a surviving mutant is statistically more likely to present virulence-increasing traits.

**Figure 7:**
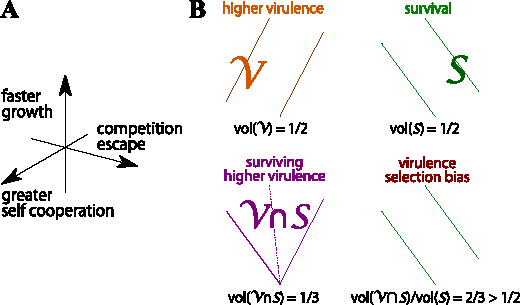
Geometrical explanation of the virulence selection bias. Leaving aside the competition pressure undergone by the resident, which does not qualitatively affect the final result, we can project the space of unconstrained mutations (scenario #3) into an arbitrarily bound cube. Each axis corresponds to the mutational deviation of a within-host trait value, as depicted on the diagram **A**. More precisely, i) the competition undergone by the mutant with itself decreases along the *x*-axis (*m*_2,2_ – *m*_1,1_) – or equivalently the self cooperation increases, ii) the competition undergone by the mutant from its ancestor decreases along the *y*-axis (*m*_2,1_ – *m*_1,1_) and iii) its the intrinsic growth rate increases along the *z*-axis (*ϱ*_2_ – *ϱ*_1_). **B)** Mutants with higher virulence than their ancestor grow faster or compete less with themselves (region *V*), but only mutants that grow faster or experience less competition from their ancestor survive (region *S*). Each of these two regions occupies half of the volume (vol) of the cube. Their intersection *𝒱* ∩ *𝒮* occupies ⅓ of the cube, which represents ⅔ of the volume of *𝒮*. Therefore, more than half of the surviving mutants are more virulent than their ancestor. Note that this explanation is also easily applicable to growth-driven (scenario #1) and competition-driven mutations (#2) by projecting over the vertical axis or the middle horizontal plane respectively. Traded-off mutations (scenario #4) cannot however be looked at this way.

### 3.2 Large mutations

Numerical simulations under Gaussian mutational distribution and the free mutational scenario not only confirm the previous results (on the left hand side of the graphs in **Figure 8A and B**), but they also provide insight to the consequences of larger mutation effects over the epidemiology and evolution of the parasite (on the right hand side).

**Figure 8:**
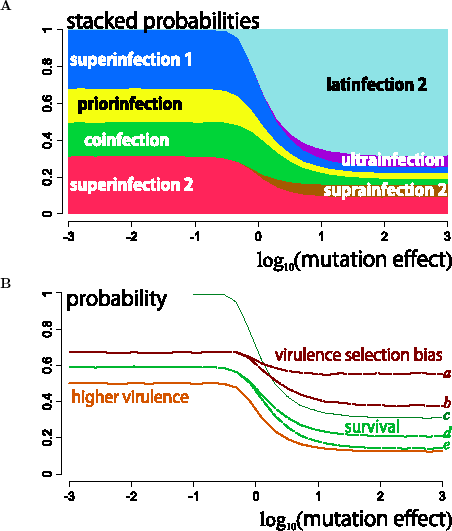
Mutant virulence and survival under large mutational effects. These results are based on numerical simulations under the following mutational setting: the mutational distributions are assumed Gaussian and follow the free mutation scenario*, i.e.* no correlation between within-host growth traits and equal standard deviation across traits, here called the mutational effect. For 30 mutational effect values ranging from 10*^—^*3 to 10^3^, we simulated 10^5^ independent (and separate) mutations of the same resident (with no loss of generality). For each of these 3*·*10^6^ mutations, we registered the induced infection pattern and compared its virulence to that of the resident. In both panels, the *x*-axis corresponds to the order of magnitude of the mutational effect, while the *y*-axis corresponds to probabilities, either stacked (panel **A**) or independent (panel **B**). The vertical dotted line separates the domain of small mutations (on the left), from the one of large mutations (on the right), arbitrarily set at one order of magnitude below the resident‘s traits absolute values. The horizontal dotted line indicates the one half probability, *y* = 1*/*2. **A**) Probability distribution of the infection patterns induced by the mutation according to the mutational effect. **B**) Probabilities that a mutation increases virulence (orange curve), mutant survival (green curves: upper bound (*c*), high (*d*) and low (*e*) extrapolations) and virulence selection bias (brown curves: high (*a*) and low (*b*) extrapolations) as functions of the mutational effect. The solid curves (or curve portions) correspond to certain data, either ground simulation outputs or outputs combined with applicable analytical results. The dashed curve portions, lying in the large mutation side of the graph, correspond to simulation outputs combined with extrapolated analytical results from the small mutational setting. See text for more details.

First but not surprisingly, latinfection 2, suprainfection 2 and ultrainfection occur more frequently when mutations get larger (Figure 8**A**), as predicted by analytical calculations. We even remark that more than two thirds (ca. 68%) of very large mutations produce the latinfection 2 pattern. This has important consequences on mutant survival since, whatever its epidemiological rates, no mutant can survive if it undergoes latinfection. Actually, latinfection 2 is the only pattern under which we can exactly assess mutant survival under large mutations. This is the meaning of curve *c* in Figure 8**B**: it represents the upper bound of the mutant survival probability, that is the sum of the probabilities of all infection patterns but latinfection 2 as if they were all allowing for mutant survival (which is nonetheless very unlikely). We can also extrapolate the survival probability from the infection pattern distribution assuming that infection patterns resulted in or prevented mutant survival in a similar way to the small mutations scenario (*i.e.* all superinfecting, all coinfecting, half of priorinfecting and none of superinfected or ultrainfecting mutants survive (see appendices F.3 and F.5)). Because we have no basis from which to extrapolate mutant survival under suprainfection 2, we consider the two extreme cases: either all suprainfecting mutants survive (curve *d* in Figure 8**B**), or none of them survives (curve *e*). These two extrapolated boundaries suggest that mutant survival is likely to lie approximately halfway between 0 and curve *c* (ca. 0.2).

Virulence (orange curve) is a ground (*i.e.* unextrapolated) output of the simulations and interestingly, while small mutations produce equal numbers of mutants with higher virulence or lower virulence with respect to their ancestor under all investigated mutational distributions and mutational scenarios, the probability that a mutant is more virulent than its ancestor dramatically drops as mutations become larger and saturates at a relatively low value (ca. 0.13). This results from the predominance of latinfection 2 in large mutational outcomes, and marginally the appearance of suprainfection 2, the two infection patterns in which mutants are avirulent by definition.

Virulence selection bias (brown curves), that is the proportion of surviving mutants which are more virulent than their ancestor relative to the proportion of surviving mutants, is found by combining the virulence mutational outcomes to the aforementioned extrapolated survival (curve *a* excludes suprainfecting mutants from survival while curve *b* includes them). Of course, the decrease in mutant survival does not affect as much virulence selection bias since the surviving mutants might be very rare but all virulent. We nonetheless observe that the virulence selection bias decreases and steps below the one half probability line when mutation effects become very large.

## 4 Discussion

The idea that virulence could be driven by within-host competition naturally arose with the first theoretical models accounting for parasite polymorphism (Levin and Pimentel, 1981; Bremermann and Pickering, 1983; Nowak and May, 1994; May and Nowak, 1995), which is now known to be the rule of natural infections (Petney and Andrews, 1998; Read and Taylor, 2001; Balmer and Tanner, 2011). While some empirical evidence suggest an evolutionary increase of virulence due to a positive correlation with competitiveness (de Roode et al., 2005; Bell et al., 2006), several experimental systems have shown that within-host parasite polymorphism would counter-select virulence instead (Turner and Chao, 1999; Garbutt et al., 2011; Pollitt et al., 2014), as predicted by alternative models (Frank, 1992; Brown and Grenfell, 2001; Brown et al., 2002; Brown, 1999; Gardner et al., 2004). These works highlight the fact that when infection relies on a collective action (such as public goods production) then lower virulent strains are favored.

Many subsequent studies tried to find a unified answer to the evolution of virulence in a multiple infection context but they are usually based on strong assumptions or lack one essential dynamical level (either within or between host dynamics), as we detail below. Following the specifications that a relevant approach to this issue has to satisfy Cressler et al. (2015), we developed a nested model based on Sofonea et al. (2015) with explicit within-host dynamics, between-host dynamics and several possible mutational settings. We paid special attention to the generality of the possible patterns each of these processes can generate. In particular, we used the infection pattern typology exposed in a previous work (Sofonea et al., 2017), that is the coupling between inoculation challenges and parasite growth outcomes, to avoid any arbitrary assumptions on the within-host outcomes and epidemiological trade-offs.

First, we show that while quite simple within-host dynamics can generate seven infection patterns, only three of them matter if mutational effects are small. These patterns are superinfection, coinfection and priorinfection (note by the way that the well-known super/co-infection dichotomy misses here an evolutionary relevant pattern). This may explain why the four other patterns are marginal in nature (but see Sofonea et al. (2017)). Second, and importantly, we show that most of the mutants able to survive are more virulent than their ancestor. We establish the robustness of this result by examining several distributions of mutational effects, including generic distributions, and various mutational scenarios, including differential variance and unfavorable genetic constraints (competitiveness coming with a cost, as spiteful compounds for instance (Gardner et al., 2004)). We therefore conclude that if polymorphism (of within-host growth traits) can emerge, then this inevitably increases the mean virulence of the parasite species.

In addition, we observed that large mutations are mostly deleterious. This provides an *a posteriori* support to our focus on small mutations since within-host growth traits appear to be, according to the model, under a strong selective pressure which denotes important developmental constraints. Further, large mutations contribute considerably less to the emergence of more virulent mutants (about three times less under the Figure 8 setting (unshown result)) and the virulence selection bias seems to be reduced. This may be explained by the fact that the vast majority of large mutations fail to produce virulent mutants which compensates the ground virulence selection bias. This question is however left open as more specific investigations on large mutations are required.

To our knowledge, our framework is the first to capture virulence evolution in multiple infections while 1) explicitly accounting for dynamics ranging from the molecular kinetics to epidemiological persistence, thus making virulence emerge from within-host growth traits, 2) allowing for several infection patterns, single and co-transmissions simultaneously, recovery, multi-dimensional mutations and 3) not specifying any link between virulence and competitiveness nor any epidemiological trade-off, 4) addressing the evolutionary outcome through weighting all possible mutations instead of considering the fate of one more virulent mutant alone and 5) being analytically tractable.

The present model captures several infection patterns with one set of equations instead of using distinct models for each pattern (Nowak and Sigmund, 2002), or deriving them from limit cases (Mosquera and Adler, 1998), or artificially mixing them (Boldin and Diekmann, 2008). The within-host outcomes completely rely on explicit population dynamics of the parasites and not any trait defined at a higher organization level such as virulence (Levin and Pimentel, 1981; Nowak and May, 1994). In particular, parasite competitiveness results from cross interaction traits that do not affect the virulence level. A parasite type can therefore be indifferently invaded and outcompeted by a more virulent and a less virulent type (see Figure 3A). This independence of competitive ability and virulence contrasts with most models (Frank, 1992, 1994; Bonhoeffer and Nowak, 1994; May and Nowak, 1994; Nowak and May, 1994; Frank, 1996; Gandon, 1998; Boldin and Diekmann, 2008).

Here, unlike Mosquera and Adler (1998), double infections originate from actual stable within-host dynamics. Unlike in May and Nowak (1995), they may indifferently affect transmission or not while the trend on virulence evolution is not affected by the co-transmission probability, as opposed to Alizon (2013b). While cooperation is allowed through cross interactions or potential increase in overall transmission in double infections, polymorphism implies increased virulence. This contrasts with earlier studies (Brown et al., 2002; West and Buckling, 2003) in which polymorphism (or low relatedness) decreases virulence. Nonetheless it has been shown that the results of these models can be inverted by considering the between-host level which they lack (Alizon and Lion, 2011). Note however that this requires assuming some trade-offs not needed here.

One can legitimately then wonder what drives the necessity of increased virulence that emerges in our model. We highlight (see appendix F.4) that a phenotypically close mutant survives in the host population in which already circulates its ancestor if it can prevent the host it infects from secondary infections by its ancestor and/or if it can grow within hosts already infected by its ancestor (which can persist or vanish). If the mutant can only protect its host, we have priorinfection. If it can only grow in hosts infected by the resident, we have coinfection. When the mutant is able to do both, it superinfects its ancestor. Coinfection guarantees the survival of the mutant since it can grow in all hosts. In the case of priorinfection, there is only a single ‘resource’ (uninfected hosts) so the mutant will persist in the host population only if the infections it causes are epidemiologically fitter than those caused by the resident (which depends on their transmission, virulence and recovery rates). Hence, a majority of surviving mutants are either superinfecting or coinfecting their ancestor. Since these two infection patterns require the ability to grow within already infected hosts, these mutants necessarily present a faster intrinsic growth or escape their ancestor‘s competitive pressure. Faster intrinsic growth is one of the two features of higher virulence – the other being greater self cooperation. Consequently, either mutants survive exclusively because of their faster intrinsic growth and they are more virulent, either they survive because they undergo less competition from their ancestor while they present greater self cooperation or faster intrinsic growth, which makes them more virulent as well. In the end, the probability that a surviving mutant is more virulent is at least of one half (and up to one), while only half of the mutants ever produced are more virulent than their ancestor. This result holds independently of whether the parasite‘s intrinsic growth trades off with competition escape.

Coming back to the adaptive explanation of virulence, we are prone to thinking that this systematic selection bias towards higher virulence answers this question by unifying reasons 2 and 3: in multiple infections, virulence results from a mixture of selection for competitiveness – actually competition escape ability – and as a by-product of local selection for faster within host growth or greater self cooperation. As pointed out by Levin and Pimentel (1981), this induces a short-sighted evolution as more virulent mutants survive even if their traits do not lead to epidemiological optimality, thus leading to an unavoidable degree of maladaptation. It is then legitimate to wonder where this increasing virulence will lead the parasite species, *e.g*. towards a stable polymorphism asymmetrically distributed towards higher virulence or a never-ending increase of virulence. This model is unable to address these long-term dynamics, however at some point epidemiological constraints will possibly limit this increase and/or the host will possibly develop resistance or tolerance.

As a counterpart of the model‘s generality, knowing that the mutant survives does not give us any insight on the survival of the resident. We claim that this does not affect the result since, whether the ancestor survives or not, the mean virulence in the parasite population increases in any case. It is however a main perspective of future work to generalize the framework to an unbounded and varying polymorphism. This task does not seem analytically tractable but specifying the functions undetermined so far based on documented experimental systems, as recommended by Cressler et al. (2015) sounds promising. Accounting for spatial structure, host heterogeneity and host evolution would certainly modulate the present result.

## Acknowledgements

The authors thank Sébastien Lion, Eva Kisdi, Yves Dumont, and Alain Rapaport for their helpful suggestions.

## Authors’ Contributions

Conceived the project: M. T. Sofonea, S. Alizon and Y. Michalakis.

Performed the analysis: M. T. Sofonea.

Wrote the article: M. T. Sofonea, S. Alizon and Y. Michalakis.

